# NeuroCI: Continuous Integration of Neuroimaging Results Across Software Pipelines and Datasets

**DOI:** 10.1101/2022.05.30.494062

**Authors:** Jacob Sanz-Robinson, Arman Jahanpour, Natalie Phillips, Tristan Glatard, Jean-Baptiste Poline

## Abstract

—Neuroimaging study results can vary significantly depending on the datasets and processing pipelines utilized by researchers to run their analyses, contributing to reproducibility issues. These issues are compounded by the fact that there are a large variety of seemingly equivalent tools and methodologies available to researchers for processing neuroimaging data. Here we present NeuroCI, a novel software framework that allows users to evaluate the variability of their results across multiple pipelines and datasets. NeuroCI makes use of Continuous Integration (CI), a software engineering technique, to facilitate the reproducibility of computational experiments by launching a series of automated tests when code or data is added to a repository. However, unlike regular CI services, our CI-based framework uses distributed computation and storage to meet the large memory and storage requirements of neuroimaging pipelines and datasets. Moreover, the framework’s modular design enables it to continuously ingest pipelines and datasets provided by the user, and to compute and visualize results across the multiple different pipelines and datasets. This allows researchers and practitioners to quantify the variability and reliability of results in their domain across a large range of computational methods.

## I. Introduction

Result variability – or robustness – is an important concern in scientific data analysis. For instance, neuroimaging results are sensitive to many software-related methodological variations such as computational environments, data analysis pipelines, pipeline versions and parameters, and statistical models. Neuroimaging results are also sensitive to data-related variations such as study populations, sample sizes, datasets, and image acquisition parameters [Carp, 2012, Kennedy et al., 2019]. Changes in any of these methodological choices can affect the results obtained by researchers, contributing to scientific result reproducibility issues. Moreover, researchers are often faced with multiple choices of pipelines featuring similar capabilities, and which may yield different results when applied to the same data. Given the large number of unique analysis procedures that can be used to carry out a single neuroimaging experiment, it is often unclear what approach neuroscientists should take when facing analytical flexibility.

In computational experiments, Continuous Integration (CI) has been proposed as a method to evaluate and facilitate reproducibility [Krafczyk et al., 2019]. CI is a technique used in software engineering whereby a series of tests is launched as soon as new code has been committed to a code repository to check if the expected software functionality is obtained. However, applying CI to a neuroimaging data analysis context involves a series of challenges due to the large storage requirements, privacy conditions of datasets, and the large memory requirements of the processing pipelines. To solve these limitations, additional software infrastructure is required.

The goal of this research project is to provide neuroimaging researchers with a software framework that is capable of carrying out neuroimaging experiments in a CI setting, embracing its reproducibility friendly features, while making use of third party distributed cloud computing and storage to overcome the aforementioned technical challenges. In neuroscience, new pipelines, pipeline parametrization, and new datasets emerge regularly. In order to evaluate the results of existing methods alongside new methods as they are developed and to address software variability concerns in the brain imaging field, the framework is modularly designed, allowing users to obtain results across multiple pipelines, parameterizations, and datasets that they may continuously specify at any stage of their experiment. This grants researchers the ability to continuously and systematically quantify and evaluate the variability and robustness of their results, and therefore have a greater degree of trust in the conclusions they draw.

More specifically, we present a scientific evaluation framework that encapsulates the following contributions to computational reproducibility and e-Science literature:

- Modular system design: allows the iteration of multiple pipelines on multiple datasets for reproducibility purposes.
- Distributed computation: Accounts for the privacy, data, and memory intensive needs of neuroimaging pipelines on large datasets, which are beyond those of regular CI services.
- Transparent workflows: Publicly keeps track of how each result is produced and stored. Up-to-date interactive visualizations of all of the newest results obtained.
- Open source, version-controlled code, available at: https://github.com/neurodatascience/NeuroCI

In addition, we present a concrete use-case for this technology, demonstrating its functionality by evaluating the variability of hippocampal volumes obtained with FSL and FreeSurfer

–two of the most commonly used neuroimaging pipelines

–on the Prevent-AD Alzheimer’s disease dataset [Tremblay-Mercier et al., 2021].

### II. Related Work

### A. Neuroimaging Reproducibility Crisis

There are multiple studies that address the effect of different datasets, pipelines, and parameters available to neuroimaging researchers on result reproducibility. One study submitted a single event-related fMRI experiment to 6,912 unique – but relevant – analysis procedures and found substantial result variability [Carp, 2012]. It emphasizes that analytic flexibility combined with selective analysis reporting could result in high false positive rates in neuroimaging research. Bhagwat et al. investigated the impact of pipeline selection on cortical thickness measures using multiple tools (ANTs, CIVET, FreeSurfer), cortical parcellations, and quality control methods on two datasets and concluded that these factors all significantly affect the results, and call for the “*need for more rigorous scientific workflows and accessible informatics resources to replicate and compare preprocessing pipelines*” [Bhagwat et al., 2021]. Another study compared automated MRI brain segmentation in FSL and FreeSurfer and found a low correlation between the segmentation results of the two tools [Quilis-Sancho et al., 2020]. Data from three task-fMRI studies that were reanalyzed using three pipelines (AFNI, FSL, and SPM) yielded marked differences when comparing the statistical maps they produced [Bowring et al., 2019]. Another study used the same three pipelines on fMRI data from healthy controls, and obtained false-positive rates of up to 70%, calling into question the validity of fMRI studies, and concluding that the pipelines may have a large impact “*on the interpretation of weakly significant neuroimaging results*” [Eklund et al., 2016]. Recently, reproducibility concerns have provoked large-scale neuroimaging multiverse analysis studies that use “*several analysis pipelines, and preferably by more than one research team*” [Botvinik-Nezer et al., 2020]. The ‘Neuroimaging Analysis Replication and Prediction Study’ provided the same neuroimaging data to 70 independent teams to analyze. No two teams chose identical analysis workflows, only one of nine hypotheses showed a high rate of significant findings, and only three hypotheses showed “*consistent non-significant findings*” [Botvinik-Nezer et al., 2020]. Another multiverse analysis on diffusion MRI fiber tractography gave 42 independent teams the same whole-brain streamline data to segment. Different segmentation protocols resulted in different reconstructions of the white matter pathways [Schilling et al., 2021]. The high variability in the results of both studies highlights the need for performing and openly disseminating multiple analyses on the same data. Pipelines are also affected by computational conditions, such as OS versions, hardware, and parallelization parameters. These numerical errors are propagated at each step of a pipeline, and can lead to error amplification [Kiar et al., 2020, Salari et al., 2020]. Additionally, available tools may have high technical proficiency requirements, which can result in suboptimal usage by researchers [Sherif et al., 2014]. The current format of academic journals, which are predominantly human-readable, can lend itself to transparency and efficiency issues when details such as parameters and software versions are not specified, or machine-readable code or data is not publicly provided, making publications challenging to reproduce [Kennedy et al., 2019].

### B. Reproducibility and Continuous Integration (CI) in Computational Science

CI has been brought up in a scientific context as a method for ensuring the reproducibility of software results across computational environments and code changes. One study points out that by using CI the authors are able to rerun an experiment “*whenever updates or improvements are made to source code or data*”. They describe benefits such as the strategy’s reliance on transparent code and accessible data facilitating Open Science, and the use of containers allowing the user to specify computational environments [Beaulieu-Jones and Greene, 2017]. Another similar study mentions that CI provides a dedicated and contained platform to find and verify data and code used to generate results [Krafczyk et al., 2019]. Scientists can use it to recreate CI-based experiments with minimal effort from within their browser. Currently, the ReproNim TestKraken [ReproNim, 2021] software is being developed to test workflows in a matrix of parametrized environments. It offers CI runs to execute tests, and allows the user to fully specify the computational environment. Overall, while CI has already been introduced in the context of scientific testing, current implementations are few, still in early development, and have not yet been comprehensively tested.

### C. Limitations of Existing CI-based Reproducibility Systems in a Neuroimaging Context

The scientific CI solutions described in the previous section have three main shortcomings in the context of neuroimaging data that limit their utility. The first is they do not feature a strategy for dealing with the large amounts of data found in neuroimaging datasets, and the large memory requirements of neuroimaging pipelines. Neuroimaging datasets often occupy hundreds of GB of disk space. Free CI services typically offer a fixed amount of processor-time or credits per week or month, and this processor time is insufficient to run the substantial computations required for neuroimaging experiments, which is better suited to HPC (High Performance Computing) clusters [Beaulieu-Jones and Greene, 2017, Krafczyk et al., 2019]. TestKraken, which is still in development, offers users who are technically competent the possibility of distributing tasks using Slurm. Users, however, are not shielded from the internals of HPCs, and would have to deal with workflow engines, schedulers, and other complexities. The second limitation of the two published studies is that their code and CI setup is geared towards running a single experiment with hardcoded processing scripts and tests for single datasets. As such it is not easy to use their code as a framework that generalizes to other computational experiments, and users would have to start from scratch in creating their own CI analyses. A generalized, modular result testing CI framework lowers the barrier of adoption for users. The third limitation of these studies is that clinical datasets often have restrictions on the data and granularity of the results the user is allowed to share or store in a public repository. They don’t have mechanisms or storage solutions that allow individual dataset usage policies to be respected.

## III. Architecture

NeuroCI consists of a collection of Python scripts and modules that are executed during CI runs and are configured by an experiment definition. The scripts are executed whenever there is an update in the GitHub repository containing the experiment definition, and can also be configured to run at regular intervals using cron scheduling. The scripts make use of the CBRAIN [Sherif et al., 2014] distributed computation system’s API to control the workflow for submitting tasks. As such, the CI framework continuously calculates and updates results, allowing the user to re-evaluate the validity of scientific claims as they add new data, processing pipelines, or parameters. In doing so it provides them with a dynamic view of result variability as new results are computed and made available to the framework. The framework uses CircleCI [CircleCI, 2022], a Continuous Integration service with a free plan that has been simple and reliable to set-up and use, and leverages many existing tools from the neuroimaging open-science ecosystem (summarized in Table 1).

**TABLE 1.** A SUMMARY OF THE DIFFERENT TECHNOLOGIES THAT ARE USED IN NeuroCI, THE Architecture SUBSECTION THAT THEY ARE MENTIONED IN, AND THEIR ROLE.

| Architecture Subsection | Role | Technology |
| --- | --- | --- |
| A | Distributed storage management | DataLad |
|  | Distributed computation management | CBRAIN |
|  | Portable pipeline containerization | Boutiques |
| B, C, D, G, H. | Transparent, open code and experiments | GitHub, Python |
| E | Continuous Integration | CircleCI |
| F | Interactive visualization | NeRV |

### A. Distributed Storage and Computation

The Continuous Testing framework relies on an ecosystem of existing tools to successfully distribute the storage and computation needs of an experiment. There are four main modules in the ecosystem, illustrated in Fig. 1. First, The ‘Distributed Computation’ module refers to CBRAIN. CBRAIN is a distributed cloud computing platform that features an API. The API allows NeuroCI to launch the heavy distributed computations on Compute Canada servers from within CircleCI runs. The particular instance of CBRAIN we will use serves over 1400 users spread across 296 cities in 44 countries as of May 2022, and is “*hosted on nine national research HPC centers in Canada, one in Korea, one in Germany, and several local research servers*” [Sherif et al., 2014]. Second, the Continuous Integration module refers to the code executing in CircleCI that orchestrates the automated workflows for experiment execution. This module is described in detail in the following subsections of the Architecture. Third, the ‘Data Retrieval’ module aims at making all of the data available in the CBRAIN distributed computation system. By default this module relies on the Datalad [Halchenko et al., 2021] framework to acquire neuroimaging data. Datalad is an open-source distributed data versioning system, which can reference datasets indexed in the Canadian Open Neuroscience Platform [CONP, 2022] (CONP) repositories. The CONP is an organization that provides infrastructure for promoting data sharing and open-science workflows. The distributed CONP data is downloaded using DataLad on Compute Canada servers, and transferred to a data provider directory where it can be accessed by CBRAIN. Template scripts are provided to automate this process. Other repositories and download methods can also be used to acquire the data if necessary. Fourth, the ‘Pipeline Containerization’ module integrates processing pipelines in CBRAIN, which is done by creating Boutiques [Glatard et al., 2018] container descriptors for them. Boutiques is a tool that facilitates the integration and execution of applications across computational platforms, and its descriptors allow the user to use the described tools on CBRAIN as well as dozens of other virtual research platforms. The descriptors are sent to the CBRAIN administrators, who integrate them after verifying that they are secure.

**Fig. 1.**
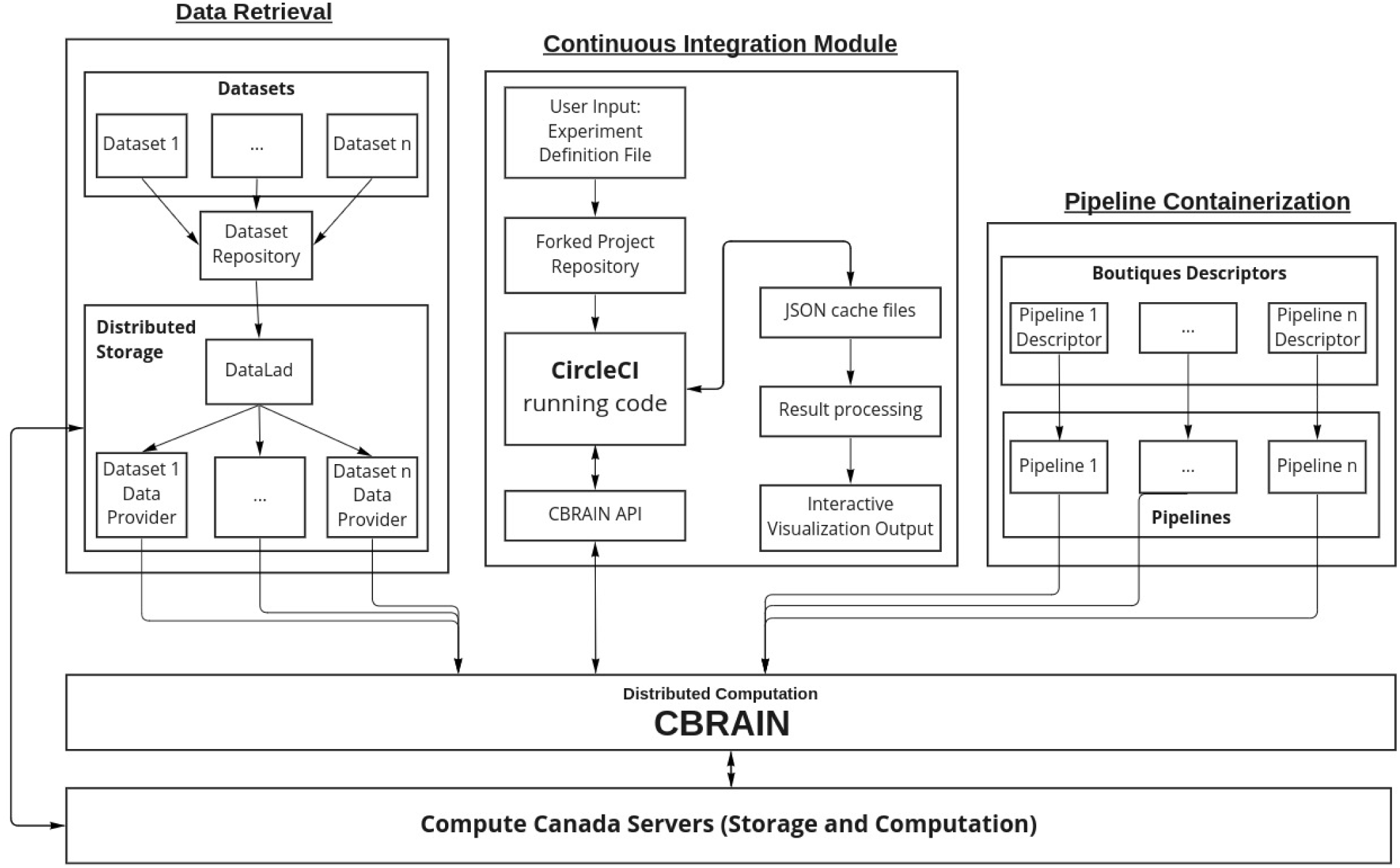
The four main modules comprising the software ecosystem of NeuroCI. Distributed computation occurs on the CBRAIN platform. The data retrieval and pipeline containerization modules make the data and tools available on CBRAIN, and the Continuous Integration module orchestrates the workflows.

### B. User Input

An ‘experiment definition’ json format file located in the working directory of the project allows the user to specify the processing pipelines, the associated parameter file paths, and datasets to obtain results for. It also allows users to specify tasks to resubmit and a blocklist of files to ignore. The ‘Config_Files’ directory contains a yaml configuration file where the user can provide the CBRAIN IDs for the data providers where the datasets are located, and the CBRAIN IDs for the pipelines. The ‘Task_Parameters’ directory contains json files that store the CBRAIN parameters for the pipelines. Lastly, the user must enter their CBRAIN credentials and CircleCI API token in the environment variables of CircleCI, which are encrypted and are unavailable to CircleCI employees. The ‘.circleci’ directory contains a configuration yaml file where the execution of the CI run is defined. It includes the software library dependencies to use, and script execution commands. We do not suggest non-advanced users modify these sections. It also includes a cron scheduling field, so users can specify the frequency with which the framework is executed.

### C. The Main Testing Loop

The main workflow and high-level logic of the CI framework is orchestrated in the ‘NeuroCI.py’ Python script. Each time a CI run is performed, CircleCI automatically uses a user-specified prebuilt Docker image (we chose Ubuntu 20.04.4 with Python 3.6.9) to build the GitHub code. We note that the distributed computations are performed on the hardware and operating system CBRAIN runs on, not in this prebuilt Docker image that orchestrates the computations. Inside the container the required dependencies are installed, and the ‘NeuroCI.py’ script is executed. The newest version of the Experiment Definition and configuration files from the repository are read, and the user is logged in to CBRAIN using its API. The script then executes the ‘main’ function which consists of a nested loop iterating across all of the datasets and pipelines listed in the Experiment Definition.

Firstly, for each dataset, the main function downloads the cache file from the previous CI run’s output artifacts, which allows the framework to access the latest task-completion information. The output artifacts of the previous CI run are retrieved using the CircleCI API, and the complete current list of tasks (and associated information) of the CBRAIN user is also fetched. The contents of the cache file are updated to reflect these newest CBRAIN task statuses. If the cache file is corrupt, nonexistent, or empty, a new cache file is created and CBRAIN is queried through the API so the cache can be populated with the relevant information (task IDs, file IDs, and completion statuses). In the case of a corrupted cache file, this avoids recomputing all the submitted tasks. After this, inside the first loop, we execute a nested loop across all of the Experiment Definition’s pipelines. For each pipeline, the current dataset’s cache file is updated in case there are any new files made available to process. The pipeline manager function is called to post tasks to CBRAIN, orchestrating the process of submitting new tasks. Finally, outside the nested loop, for each dataset, CBRAIN is queried for completed task results, and the cache file is populated with these results. The detailed implementations of these operations and files will be discussed in the following sections.

### D. Task Provenance Cache Files

Provenance of the tasks that are posted on CBRAIN is kept track of in a series of json format ‘cache’ files. One cache file is created for every dataset specified in the experiment definition. Every file inside the dataset’s Compute Canada data provider directory has a corresponding entry in the cache file and under this entry there are fields for each of the pipelines that are specified in the experiment definition. These pipeline fields are populated to keep track of the CBRAIN IDs of the tasks, inputs, and outputs, as well as the completion status for every component in the pipeline chain. We note that the cache keeps tracks of the IDs to locate the files and tasks located on CBRAIN, and not the files or tasks themselves. Since the components of the pipeline are executed sequentially, new tasks are only posted once the previous task in the pipeline has finished, and the completion is marked in the cache file. At the end of the CI run, these cache files are exported as artifacts which can be downloaded through the CircleCI API and updated in the next CI run.

### E. Continuous Integration Framework

The ‘cacheOps.py’ file is imported as a module in the main testing loop, and contains all of the functions that create, modify, or interact with these cache files. These functions perform jobs such as downloading the cache files from the last CI run’s output, creating new cache entries for newly integrated files, posting tasks to CBRAIN, and updating the cache entries when task statuses change, amongst others. The ‘cbrainAPI.py’ file is imported in ‘cacheOps.py’ and contains functions for all of the API calls made to CBRAIN. The visualization modules also use the API calls along with the output cache files to generate a summary of the experiment’s computations for the CI run. This software architecture can be seen in Fig. 2.

**Fig. 2.**
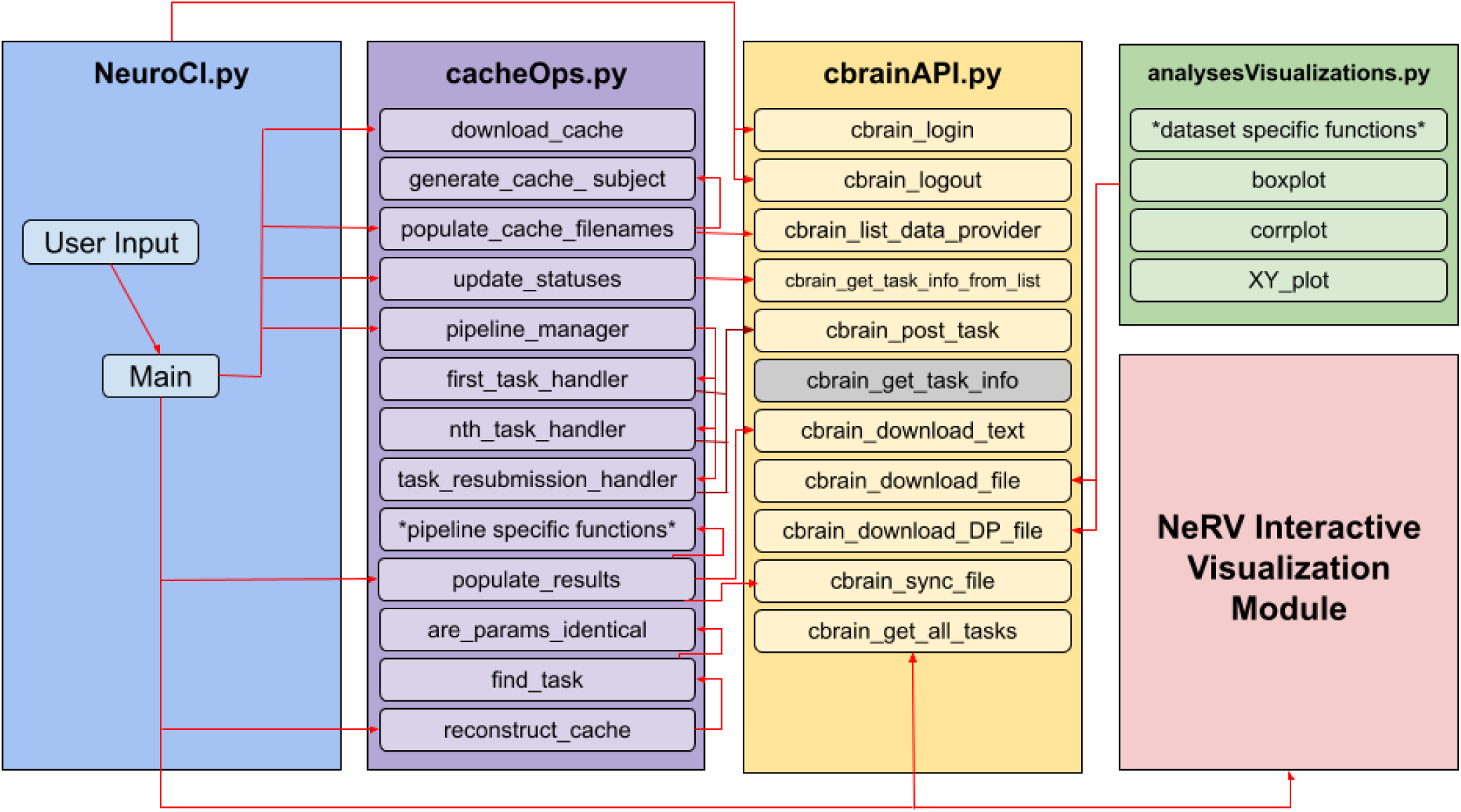
The software architecture of NeuroCI. The main testing loop is initiated in NeuroCI.py. It uses cacheOps.py to orchestrate the workflows through a series of cache files, and cbrainAPI.py to interact with the distributed computation system. Results obtained can be visualized through analysesVisualizations.py and the NeRV interactive visualization module.

### F. Interactive Visualization

NeuroCI makes use of the purpose-built Neuroimaging Results Visualization (NeRV) Python library to interactively visualize computed results. NeRV is an open-source interface developed using Plotly Dash and Pandas, and can be found at https://github.com/rmanaem/nerv. NeRV takes as input the cache files generated by NeuroCI, and generates a GUI with two tabs; one for distribution histograms, and one for joint scatter/distribution plots based on each pipeline-dataset combination. Both tabs feature interactive plots and a dynamic information panel with summary statistics relating to the computations, such as the total number of null and valued data points, and both plots feature a slider on the x-axis that allows the user to set the domain. When a CI run ends, the generated cache files are transferred automatically to a directory on Compute Canada where the NeRV interface is wrapped with a flask application and served using a WSGI entry point and Nginx server on a virtual machine running Ubuntu 20.04. The server returns a link in the CI run console for the user to visualize results in their browser.

### G. Phenotypic/Behavioral Data Integration

Researchers will often want to study the relationship between the results they computed using the framework and behavioral or phenotypic data from the datasets. This data is often registered or private, requiring subject anonymity. Coupled with the wide variety of processing and visualization methods researchers may wish to employ and the varying formats of the input data, this is beyond the scope of the framework’s interactive visualization tool. Instead, we opt to allow researchers to include their own analyses and visualizations in the analysesVisualizations.py module. The phenotypic data CSV file is downloaded from CBRAIN, using the user’s credentials for authentication in the API. The required processing is carried out to obtain the relevant age information, and a static, anonymized graph is generated using Matplotlib. The graph image file is then outputted as an artifact of the CI run. When the CI run ends, the data expires along with the container it was downloaded into, so it is not accessible to the public.

### H. Outlier Removal

We have provided two mechanisms for outlier removal in the analysesVisualizations.py module. The first mechanism consists of a simple upper and lower bound filter computed as a percentile of the distribution, where values above or below these bounds are removed from the calculations and visualizations. The second mechanism consists of a discrepancy filter where users can provide a discrepancy threshold parameter. If the discrepancy for a given observation between any two pipelines is larger than the user-provided threshold, then the data point is excluded from calculations and visualizations. This doubles as a useful method for researchers to perform quality control on the results they obtain by comparing them to the results of different pipelines.

## IV. Use-Case Results

### A. Hippocampal volume variability

The hippocampus is a structure that is commonly associated with memory function and dementia. In particular we examine left hippocampal volumes as multiple studies propose that the left hippocampus experiences a larger decrease in volume (and at a faster rate) than the right hippocampus. Multiple studies propose it has a higher discriminative power for identifying the progression of Alzheimer’s disease than the right hippocampus, and that it is a promising surrogate marker of the disease progression [Schuff et al., 2009, Vijayakumar and Vijayakumar, 2013].

However the degree and rate of left hippocampal volume loss the studies report varies with the segmentation method and datasets utilized. This is particularly evident when automated segmentation pipelines such as FSL and FreeSurfer are used [de Flores et al., 2015, Leung et al., 2010, Suppa et al., 2016]. It is unclear which pipeline’s result will be the best biomarker. As such, executing multiple pipelines on a longitudinal Alzheimer’s progression dataset like Prevent-AD is a promising starting point to ensure researchers are obtaining reliable measurements with automated segmentation methods on this important potential biomarker.

The FSL 5.0.9 [Patenaude et al., 2011] and FreeSurfer 6.0.0 [Fischl, 2012] pipelines were integrated in the NeuroCI framework, and used to find the left hippocampal volume of subjects from the Prevent-AD dataset. The Prevent-AD registered dataset is comprised of 308 cognitively healthy participants between the ages of 55-88 years old that are at risk of developing Alzheimer Disease. The participants, of which 70% are female, were followed longitudinally in multiple visits since 2011, and multimodal measurements were obtained at each visit, including the anatomical T1-weighted MRI scans that were used to obtain the following results [Tremblay-Mercier et al., 2021]. FSL produced some volumes which seem equivocally large or small, for example, results under 1500mm^3^ (11th percentile of the results obtained) or above 6000mm^3^ (99th percentile) [Honeycutt and Smith, 1995]. FSL produced a larger range of values, and the distribution of its results is asymmetrical; the results are skewed toward lower volumes than those of FreeSurfer. FreeSurfer’s result distribution is roughly symmetrical. This can be observed in the NeRV visualization output from Fig. 3.

**Fig. 3.**
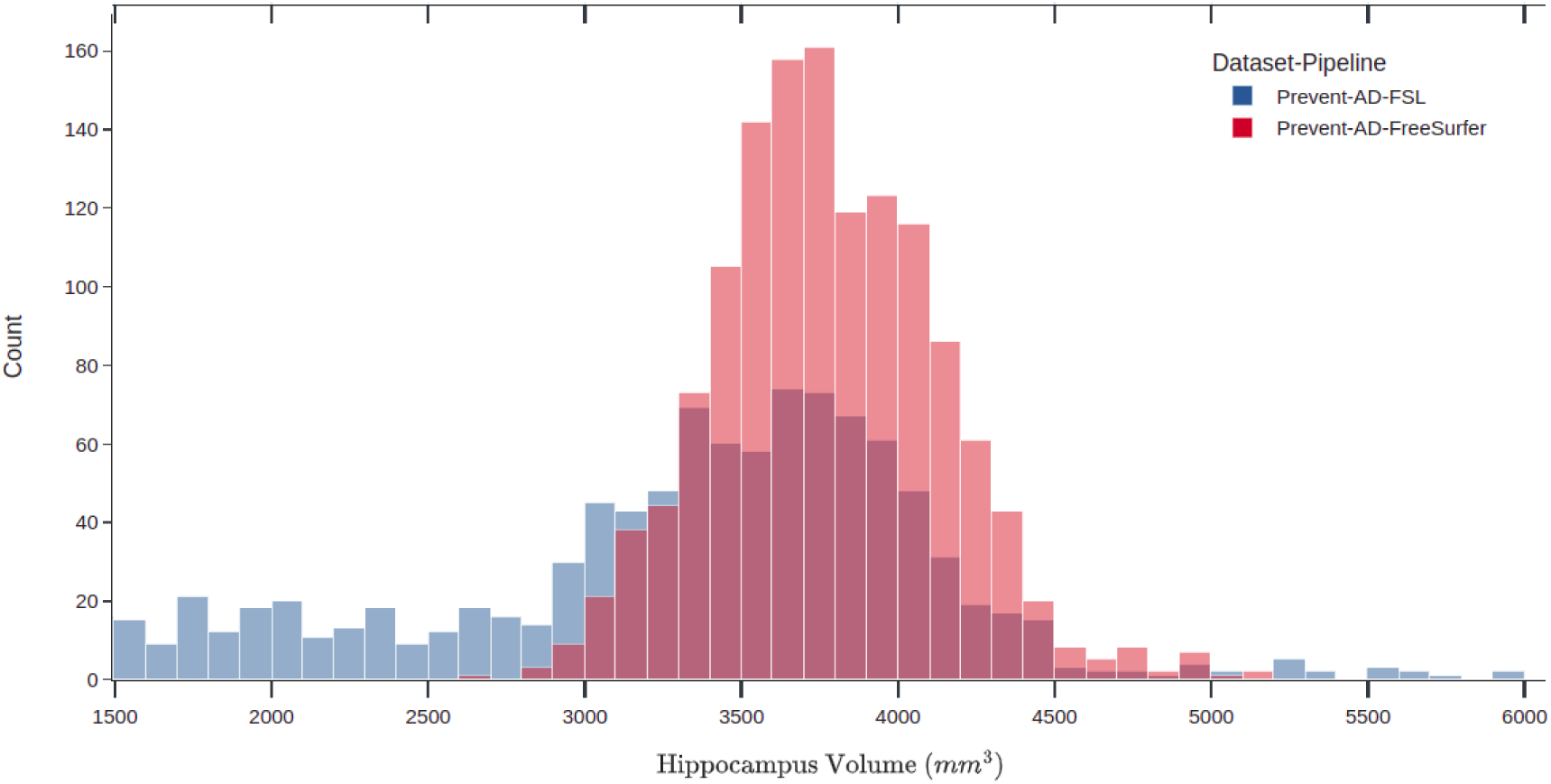
Distribution of hippocampal volumes results for FSL and FreeSurfer pipelines on Prevent-AD data after outlier values under 1500mm^3^ or above 6000mm^3^ are removed, as seen in the NeRV interactive visualization. FSL’s distribution is more asymmetrical, being skewed towards lower values.

We generated XY plots showing the correlation between the hippocampal volumes generated by the FSL and FreeSurfer pipelines. These plots can be seen in Fig. 4, and show the results before outlier removal, using bounded outlier removal, and using inter-pipeline discrepancy outlier removal. From the possible 1392 possible data points, 111 were removed due to failed segmentations across one or both pipelines. FreeSurfer consistently produced higher volumes than FSL. Before outlier removal, 1281 data points were plotted and the Pearson correlation coefficient between the hippocampal volumes generated by the FSL and FreeSurfer pipelines was of 0.0429. After bounded outlier removal (results under 1500mm^3^ or above 6000mm^3^), 987 of the data points remained, and yielded an inter-pipeline correlation of 0.2003. After inter-pipeline discrepancy outlier removal, using an inter-pipeline discrepancy threshold of 450mm^3^, 636 data points remained, and yielded a correlation of 0.8406. Scans with a very large hippocampal volume discrepancy between the two pipelines occurred in cases where FSL produced clear segmentation failures. Two examples of such cases flagged by the pipeline discrepancy outlier detection mechanism can be seen in examples 1 and 2 in Fig. 5. Examples 3 and 4 in the same figure, however, illustrate that significant pipeline discrepancies can also occur in successful segmentations.

**Fig. 4.**
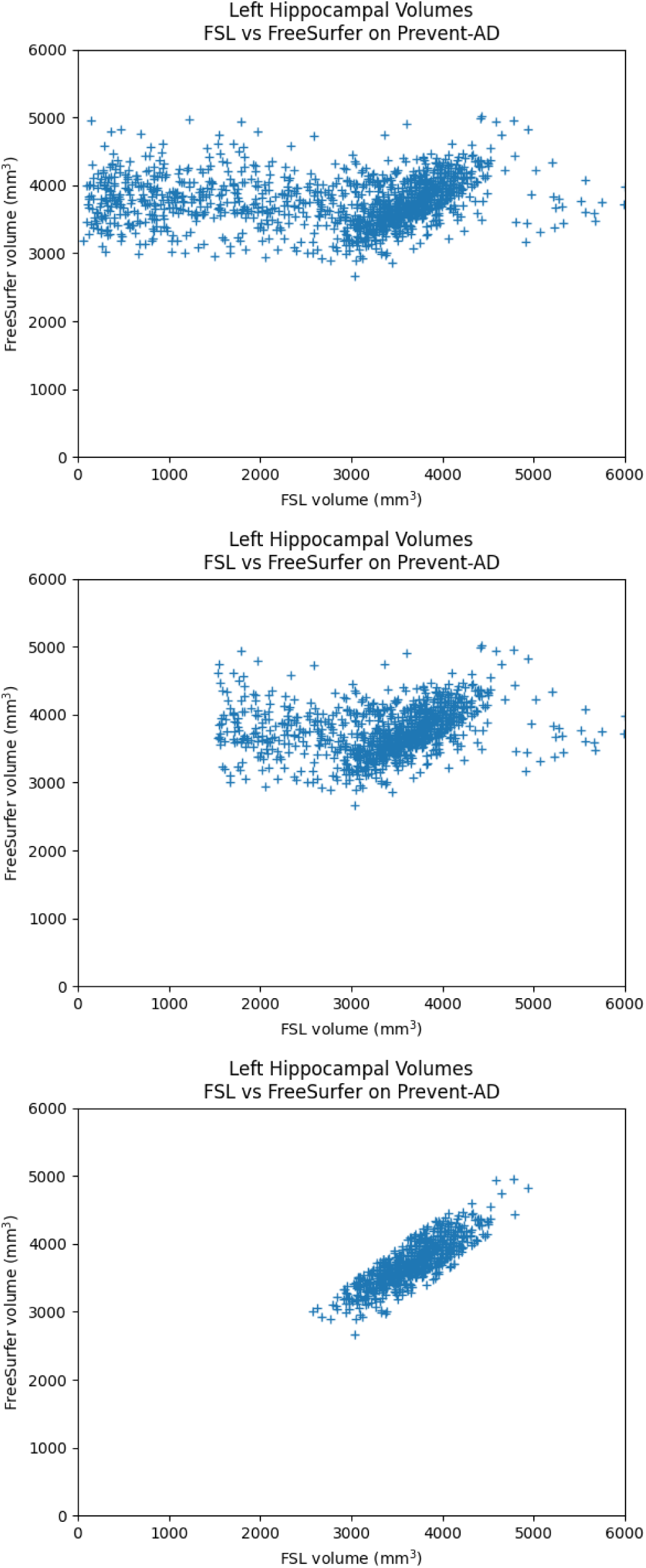
XY plots showing the correlation between the left hippocampal volumes obtained on Prevent-AD data by the FSL and FreeSurfer pipelines. FreeSurfer consistently produced higher volumes than FSL. (Top: No outliers removed, Middle: Outliers under 1500mm^3^ or above 6000mm^3^ removed, Bottom: Results with an inter-pipeline discrepancy larger than 450mm^3^ removed.)

**Fig. 5.**
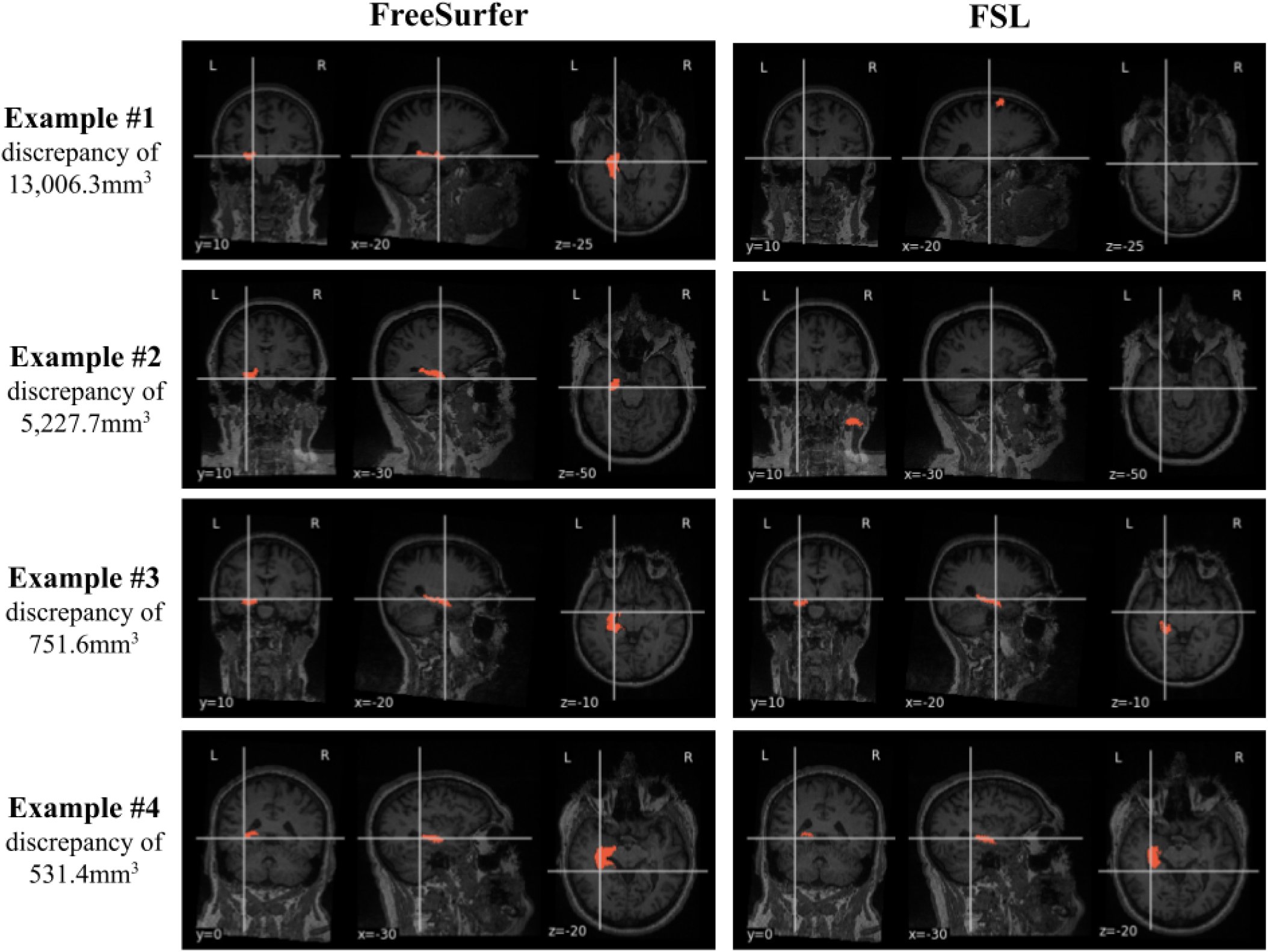
Examples of hippocampal segmentations that were flagged as outliers using FSL and FreeSurfer volume disagreements. Examples 1 and 2 have very large discrepancies due to FSL segmentation failures. Examples 3 and 4 show significant pipeline discrepancies occurring in successful segmentations that produce biologically plausible volumetric measures.

### B. Hippocampal Volumes and Age

We have included a sample workflow in the analysesVisualizations.py module to demonstrate the Phenotypic/Behavioral Data Integration and Outlier Removal, showing the relationship between left hippocampal volume and age for FSL and FreeSurfer pipelines on the Prevent-AD data. We show how this relationship varies when we apply NeuroCI’s two outlier removal strategies. Fig. 6 shows the relationship without the removal of any outliers, based on 1302 data points for FSL and 1356 data points for FreeSurfer. Fig 7. shows the relationship when values under 1500mm^3^ or above 6000mm^3^ removed, based on 993 remaining FSL data points, and the same 1356 FreeSurfer data points (no outliers were removed using this method). Fig. 8 shows this relationship when results that differ by more than 450mm^3^ are removed, and is based on 636 data points for both pipelines due to the symmetry of the removal process.

**Fig. 6.**
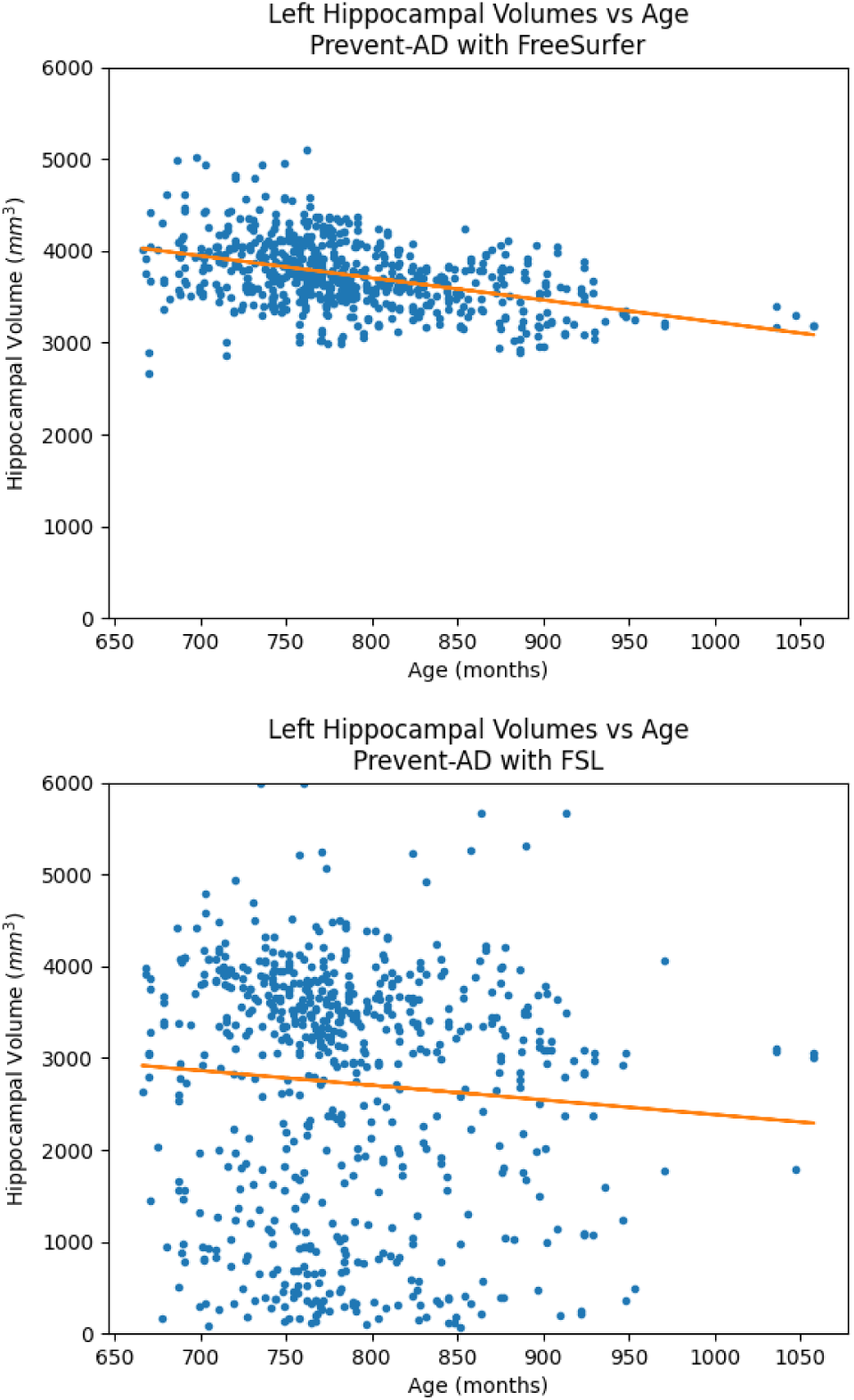
Left hippocampal volumes (Top: FreeSurfer, Bottom: FSL) plotted against subject age for Prevent-AD data without the removal of any outliers.

**Fig. 7.**
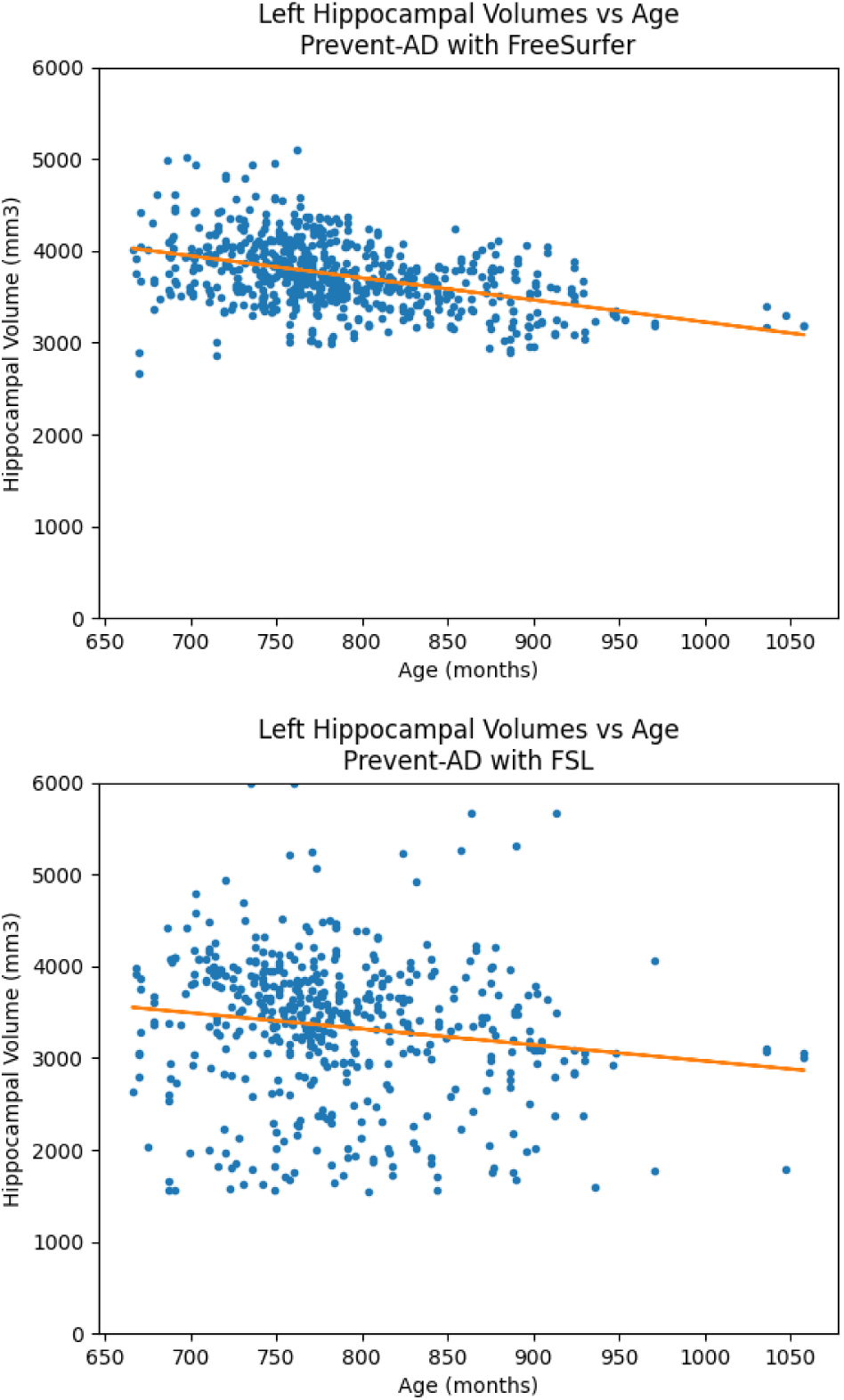
Left hippocampal volumes (Top: FreeSurfer, Bottom: FSL) plotted against subject age for Prevent-AD data. Bounded outlier removal was applied: data points under 1500mm^3^ or above 6000mm^3^ were removed. Notice that this method resulted in no outliers removed for FreeSurfer.

**Fig. 8.**
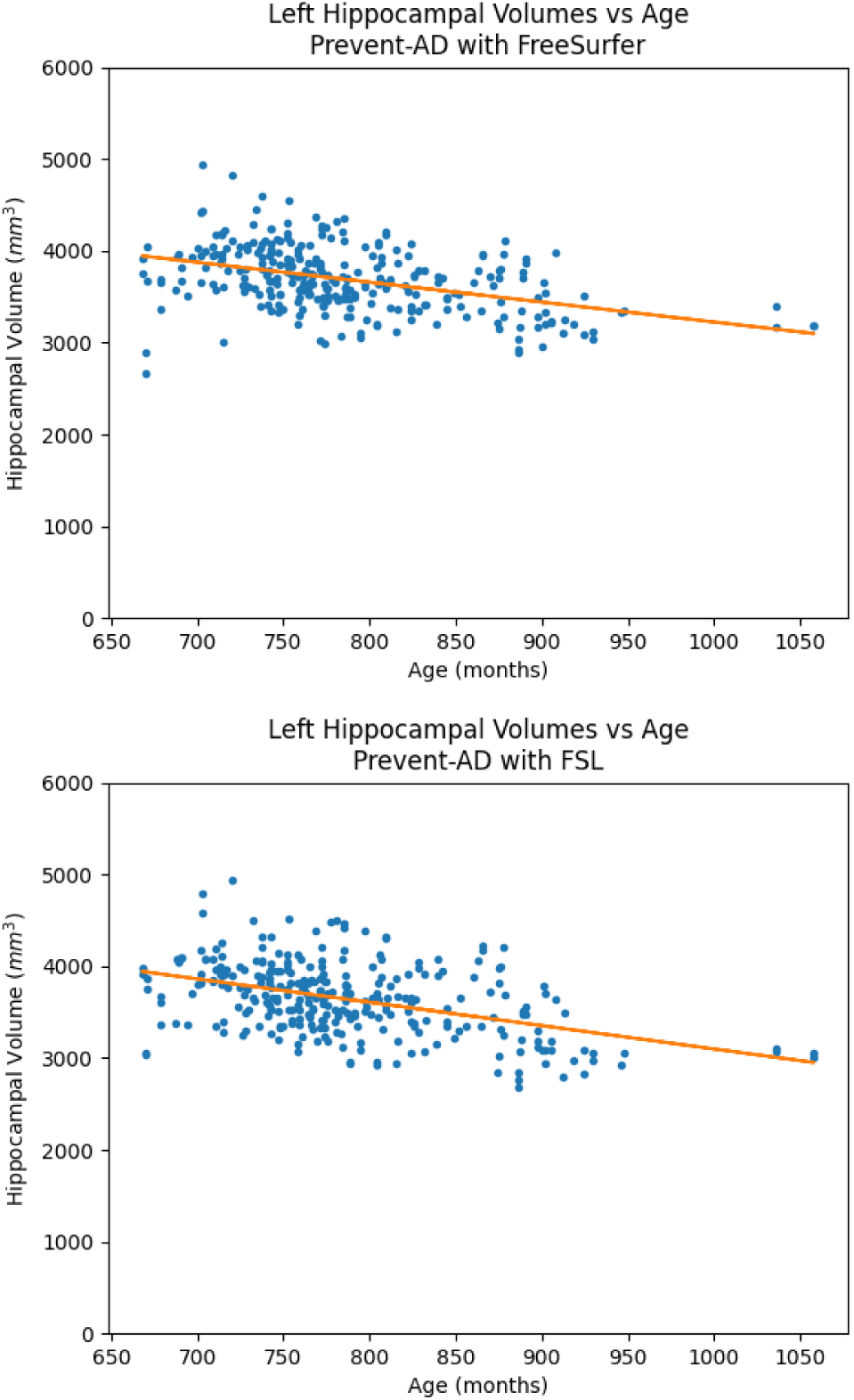
Left hippocampal volumes (Top: FreeSurfer, Bottom: FSL) plotted against subject age for Prevent-AD data. Discrepancy outlier removal was applied: results that differed by more than 450mm^3^ between the two pipelines were removed.

## V. Discussion

The limited agreement between the results of the two pipelines in the previous section illustrates the importance of a system capable of evaluating the reproducibility of neuroimaging experiment results. These results are consistent with past literature, in particular the lack of correlation in both the FSL and FreeSurfer pipeline volumetric outputs, and the bias of FreeSurfer consistently producing higher volumes than FSL are reproduced [Quilis-Sancho et al., 2020]. The lack of a ground truth between the different software methods motivates further use of the framework to properly understand the variability of results in the field, and the need to integrate more pipelines and datasets to investigate the robustness of volumetric studies in neurodegenerative pathology neuroimaging literature. This will be a next step in our future work.

Different research teams may define the hippocampus differently in terms of its constituent anatomical substructures, which can affect volumetric study results. For both FSL and FreeSurfer the anatomy of the training data was was defined and manually segmented as per the Center for Morphometric Analysis protocol from the Massachusetts General Hospital. In the case of the hippocampus, this consists of the hippocampal formation, which is “*comprised of the dentate gyrus, the ammonic subfields (CA1, CA2, CA3, CA4), the prosubiculum, and the subiculum*” [CMA, 2003]. Given that both tools agree on their definition and labelling of the hippocampus, it seems likely that the different algorithms or the different training data the segmentation tools employ are the cause of the result discrepancies.

While the proposed CI framework successfully automates the iterations of FSL First and FreeSurfer on Prevent-AD data, the absence of human inspection in the workflow leaves room for error. Experiment-specific quality control needs to be embedded in the framework or in the experiment’s process by the user to exclude incorrect results. In our use-case an error can take the form of an incorrect segmentation leading to inaccurate volumetric measures. Assessing segmentation quality is a challenge for which the gold standard is currently manual inspection by experts, but alongside our two proposed methods for outlier flagging and removal, further automated measures like the contrast-to-noise ratio of the segmentations, or the Dice scores obtained by overlapping the segmentations with an atlas could be implemented as a flagging mechanism for experts to evaluate suspicious results. In particular, the use of discrepancies between pipeline results to flag outliers is a novel feature that is useful in facilitating quality control in studies, and further motivates the development of the tool. It is also worth mentioning that due to the difficulty in segregating between public and private data in human studies, it was necessary to support local post-processing and visualization for private data, as opposed to building the online infrastructure necessary to authenticate such processes.

NeuroCI assumes that the output of the integrated pipelines is deterministic. If an experiment or pipeline’s parameters includes random seeds that affect the results, then NeuroCI is not equipped to perform numerous trials to account for this random variability. While a given tool can be parameterized in a different way and integrated as a separate pipeline in NeuroCI in order to obtain results for the different parameterizations, this requires manual integration of the parameters. NeuroCI was not designed as a tool for performing exhaustive pipeline parameter exploration. Given that hardware architecture and computational environment variations can significantly impact neuroimaging results, discrepancies that are present in the machines of the HPC network hosting CBRAIN could potentially limit the reproducibility of the results obtained using NeuroCI. However, since all the tools in NeuroCI are Boutiques tools and are therefore containerized with their dependencies, they don’t use any software on HPC machines other than the kernel, minimizing the potential impact of machine heterogeneity. Nevertheless, these discrepancies should be explicitly avoided when the pipelines are integrated on CBRAIN to ensure the determinism of the computed results.

The framework’s modular design permits users to specify their own pipelines, datasets, and parameters, allowing it to generalize to other neuroscience domains beyond our use-case with minimal modifications. Users can systematically evaluate the variability and robustness of results within their neuroscience domain, pinpoint result discrepancies caused by their methodological choices, and quantify the impact they have on the results. This will allow researchers to easily verify results in existing publications as well as their own research, and discover biases or trends impacting reproducibility. Computing the results of existing methods alongside new methods as they are developed can facilitate the consolidation of knowledge in the field. Similarly, software developers could make use of NeuroCI to evaluate their tools over multiple datasets, or compare the performance different versions of their tools. The framework’s main limitations in terms of barriers-to-entry are its reliance on CBRAIN API calls (users must have an account on a running instance of CBRAIN on a HPC), and the technical proficiency required to integrate their own pipelines, datasets, and configurations when required, which requires familiarity with GitHub, containers, and compute servers.

## VI. Conclusion

In this work we constructed NeuroCI, a generalized software framework for the continuous testing of neuroimaging results across multiple pipelines and datasets. The framework is built around the concept of Continuous Integration to facilitate result reproducibility, but leverages distributed storage and computation to meet the privacy, data, and memory intensive needs of neuroimaging research, which are beyond those offered by regular CI services. Modular software design grants users the ability to continuously integrate new pipelines and datasets, allowing the tool to easily generalize to many neuroimaging domains and scientific workflows. To demonstrate the tool we computed left hippocampal volumes using the popular FSL and FreeSurfer pipelines on Prevent-AD data, and observed a low correlation between the results of the two pipelines. These results, along with the detection of outliers based on inter-pipeline agreement motivates integrating more datasets and pipelines in the framework. The framework allows users to consolidate knowledge and quantify or explain the uncertainty of a specific neuroscience result. Confirming the reliability of results can impact researchers and practitioners by pointing to the most likely robust biomarkers. Ultimately, the platform could provide a framework for “living papers”, where scientific results would be updated over time based on the availability of new pipelines and datasets.

## VII. Acknowledgments

This work was partially funded by the National Institutes of Health (NIH) NIH-NIBIB P41 EB019936 (ReproNim) NIH-NIMH R01 MH083320 (CANDIShare) and NIH RF1 MH120021 (NIDM), the National Institute Of Mental Health under Award Number R01MH096906 (Neurosynth), the Canada First Research Excellence Fund, awarded to McGill University for the Healthy Brains for Healthy Lives initiative (NeuroHub) and the Brain Canada Foundation (Canadian Open Neuroscience Platform) with support from Health Canada, through the Canada Brain Research Fund in partnership with the Montreal Neurological Institute. This work was also partially funded by a a joint grant of the McConnell Brain Imaging Centre and the Concordia PERFORM Center.

